# Cerebellar deep brain stimulation as a dual-function therapeutic for restoring movement and sleep in dystonic mice

**DOI:** 10.1101/2023.10.30.564790

**Authors:** Luis E. Salazar Leon, Linda H. Kim, Roy V. Sillitoe

## Abstract

Dystonia arises with cerebellar dysfunction, which plays a key role in the emergence of multiple pathophysiological deficits that range from abnormal movements and postures to disrupted sleep. Current therapeutic interventions typically do not simultaneously address both the motor and non-motor (sleep-related) symptoms of dystonia, underscoring the necessity for a multi-functional therapeutic strategy. Deep brain stimulation (DBS) is effectively used to reduce motor symptoms in dystonia, with existing parallel evidence arguing for its potential to correct sleep disturbances. However, the simultaneous efficacy of DBS for improving sleep and motor dysfunction, specifically by targeting the cerebellum, remains underexplored. Here, we test the effect of cerebellar DBS in two genetic mouse models with dystonia that exhibit sleep defects— *Ptf1a^Cre^;Vglut2^fx/fx^*and *Pdx1^Cre^;Vglut2^fx/fx^*—which have overlapping cerebellar circuit miswiring defects but differing severity in motor phenotypes. By targeting DBS to the cerebellar fastigial and interposed nuclei, we modulated sleep dysfunction by enhancing sleep quality and timing in both models. This DBS paradigm improved wakefulness (decreased) and rapid eye movement (REM) sleep (increased) in both mutants. Additionally, the latency to reach REM sleep, a deficit observed in human dystonia patients, was reduced in both models. Cerebellar DBS also induced alterations in the electrocorticogram (ECoG) patterns that define sleep states. As expected, DBS reduced the severe dystonic twisting motor symptoms that are observed in the *Ptf1a^Cre^;Vglut2^fx/fx^* mutant mice. These findings highlight the potential for using cerebellar DBS to improve sleep and reduce motor dysfunction in dystonia and uncover its potential as a dual-effect *in vivo* therapeutic strategy.

## Introduction

Dystonia is comprised of an array of motor disorders and symptoms that are characterized by involuntary over-contractions and/or co-contractions of the muscles. The resulting changes culminate in abnormal postures and movements, which ultimately impose a considerable burden on the quality of life of affected individuals^1,2^. The pathophysiological underpinnings of dystonia are complex and remain only partially resolved. Contributing to this complexity is the multifaceted etiology of dystonia, ranging from hereditary factors to cerebral trauma; with a significant number of isolated dystonia cases being deemed idiopathic^2,3^. Central to our current understanding of dystonia’s pathology is the “dystonia network”, an intricate network of interconnected brain regions. Functional defects in this network are implicated in dystonia in both clinical cases and animal models^4,5^. Notably, within this network, the cerebellum has emerged as a pivotal node in the manifestation of dystonic behavior. Recent research in mouse models has started to elucidate the mechanisms for how the initiation of dystonia might act in response to cerebellar dysfunction^6–8^, with corroborating evidence observed in human patients^4,9^, especially within the context of the cerebello-thalamo-cortical circuit^10^. Interestingly, in addition to its prominent role in dystonia, cerebellar dysfunction has also been linked to specific impairments in sleep timing and quality^3,11,12^. Indeed, individuals with dystonia frequently experience sleep disturbances, including insomnia (indicative of a difficulty in falling asleep), parasomnias (indicative of a difficulty in staying asleep), and notably, pronounced deficits in the duration and quality of rapid eye movement (REM) sleep^13–15^. Analogous observations are reported in mouse models of dystonia, in which mutant mice exhibit an increase in awake time (driven by longer wake bouts), diminished REM duration (driven by fewer REM bouts), and increased latency to initiate REM sleep^16^. These data underscore the multifaceted role of the cerebellum in dystonia and suggest that it may have a broader neurological impact that extends beyond motor control to include the regulation of sleep.

Dystonia-associated sleep dysregulation is known to contribute to a myriad of comorbid disorders^17,18^ and it can also directly worsen the motor symptoms inherent to the disease^14,19^. Consequently, a prevailing challenge in dystonia therapeutics involves the simultaneous management of both motor and non-motor (sleep-related) symptoms. Deep brain stimulation (DBS) has been widely accepted as a viable treatment option to mitigate the motor symptoms in dystonia^20–23^. Alongside its role in addressing motor dysfunction, DBS also holds promise for the improvement of sleep disturbances, demonstrated by its potential to sustain the sleep state in which it is applied^24–27^. While the precise mechanism(s) through which DBS improves neural dysfunction remains unclear, the leading hypotheses suggest that DBS could be enhancing, silencing, or disrupting aberrant cellular signaling^28^. While the conventional targets for DBS in human dystonia patients remain mainly limited to basal ganglia and thalamic structures, cerebellar DBS is increasingly being explored not only due to the role of the cerebellum in motor disease pathology, but also its apparent efficacy as a therapeutic target in motor diseases such as dystonia and ataxia, in which the globus pallidus and sub-thalamic targets have provide limited therapeutic benefit for some patients^29^. Indeed, we previously showed that DBS targeted to the cerebellar interposed nucleus significantly reduces the severe motor symptoms in dystonic mice and promising results have been reported in hemidystonia patients that received cerebellar dentate nucleus DBS^7,20,30^.

Recent studies, including our own, propose a bidirectional role for the cerebellum in dystonia, influencing both motor and non-motor (sleep) impairments^11,16^. Given this dual function, the cerebellum emerges as an appealing target for DBS intervention to address both motor and sleep dysfunctions in dystonia. Therefore, here, we explore the potential of cerebellar nuclei DBS as a multifunctional therapeutic strategy, which could be used to manage both motor and sleep dysfunctions in dystonia. Towards this, we examined the *Ptf1a^Cre^;Vglut2^fx/fx^* and *Pdx1^Cre^;Vglut2^fx/fx^* mouse models of dystonia. We focused on the impact of modulating the cerebellar circuit with DBS on sleep and motor disturbances, while accounting for the differences in the severity of motor symptoms and cerebellar circuitry changes in each model. Each model showcases a specific severity in its overall motor phenotypes (severe dystonia in *Ptf1a^Cre^;Vglut2^fx/fx^*mice including persistent twisting postures and tremor, with no such overt and unprovoked motor symptoms in *Pdx1^Cre^;Vglut2^fx/fx^* mice). The differences in dystonic phenotypes are driven primarily by differential expression patterns of *Ptf1a* and *Pdx1* in the cerebellar circuit^7,8^, leading to shared yet different degrees of silencing the olivocerebellar input to Purkinje cells, which drives dystonic motor behaviors and is a circuit implicated in human dystonia^31^. Together, these models provide a unique platform for comparative analysis and for tracking how cerebellar DBS affects motor dysfunction and sleep; their patterns, timing, and quality of sleep were recently reported^16^.

In this work, we test whether cerebellar interposed and fastigial nuclei DBS can reduce sleep deficits in both the *Ptf1a^Cre^;Vglut2^fx/fx^* and the *Pdx1^Cre^;Vglut2^fx/fx^* mutant mice. In the absence of the stimulation, *Ptf1a^Cre^;Vglut2^fx/fx^* and *Pdx1^Cre^;Vglut2^fx/fx^*mice display increased awake time, increased NREM time, and decreased REM time. During stimulation, the mutant mice from both groups showed restored awake and REM sleep time. Cerebellar nuclei DBS also normalized REM latency in both groups of mutant mice, which was elevated prior to stimulation, similar to human patients with dystonia^13^. Spectral analysis of ECoG signals also showed that the DBS induced alterations in spectral power in frequency bands that are relevant for defining sleep states, indicative of changes in sleep physiology. Additionally, in the *Ptf1a^Cre^;Vglut2^fx/fx^* mutant mice with severe dystonic motor symptoms, DBS reduced the amplitude of EMG during all stages of sleep, indicating that motor function was normalized during sleep. Our work suggests that the cerebellum could be a potential dual therapeutic target for restoring movement and sleep, and that the application of cerebellar DBS may be a viable option to fill an existing gap in treating dystonia.

## Results

### Targeting the cerebellum with DBS does not impair the acquisition of ECoG/EMG recordings, nor the ability to classify arousal states

Previously, in a series of studies, we employed ECoG/EMG recordings to evaluate sleep quality and used cerebellar DBS to mitigate motor dysfunction in mouse models^7,16,20,30^. However, the unified application of both techniques requires validation to ensure that they would not adversely interfere with one another *in vivo*. We therefore implanted *Ptf1a^Cre^;Vglut2^fx/fx^*and *Pdx1^Cre^;Vglut2^fx/fx^* mice with prefabricated ECoG/EMG headmounts, and with twisted bipolar Tungsten DBS electrodes. The ECoG/EMG headmount was oriented such that the recording electrodes were situated above the frontal and parietal cortices, and the DBS electrode was oriented above the cerebellum (Figure 1A). DBS electrodes were targeted bilaterally to terminate between the interposed and fastigial cerebellar nuclei, due to their roles in movement-regulation^32^ and connections to sleep^33–35^, respectively. The electrodes terminated ∼2.5mm from the surface of the brain, within the cerebellar white matter just above the nuclei. (Figure 1E). Sleep was recorded continuously for 8 hours during the light phase (when mice naturally sleep) for four consecutive days, so that sleep quality could be assessed pre-DBS, during DBS, and post-DBS (Figure 1B-C). On the days when the DBS was delivered, the mice were stimulated continuously for the entire duration of the 8 hour recording period, with a 130 Hz biphasic stimulus (30 μA, 60 μs duration) similar to previous work^7^ (Figure 1D). Raw ECoG/EMG waveforms showed that the overall DBS paradigm and targeting did not affect the quality of the recording, nor did it affect the ability to classify the stages of wake, REM, or NREM sleep (Figure 1F). To confirm the integrity of the recorded data, we quantified the total number of artifacts observed across all mice in each group. Artifacts were automatically noted during offline sleep scoring (see Methods). Our analysis revealed no significant differences in the proportion of artifacts scored across recording days, ensuring the consistency and reliability of our experimental setup and data collection (Figure 1G).

**Figure 1:**
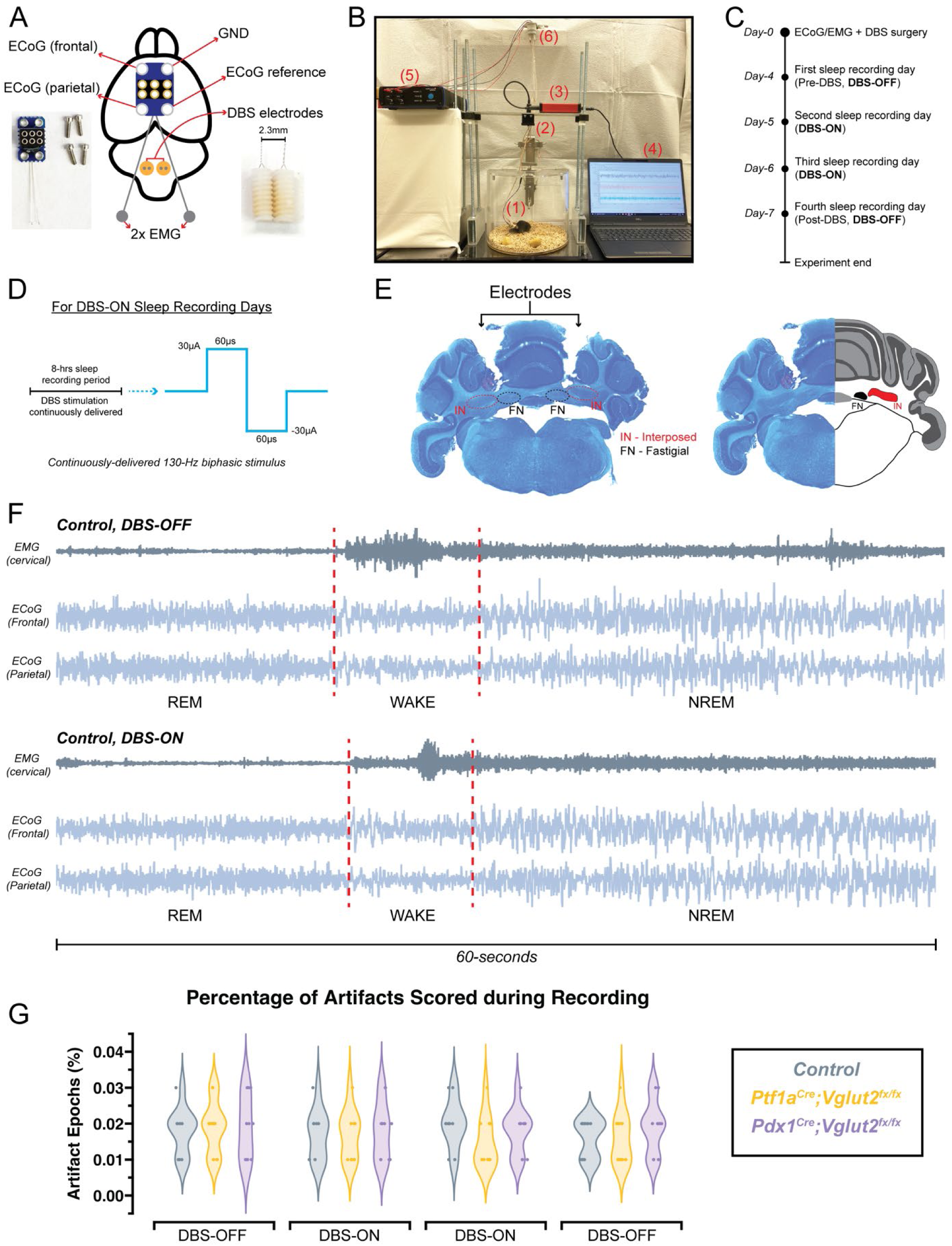
Simultaneous recordings of ECoG and EMG during cerebellar nuclei DBS. **(a)** A schematic illustration of a mouse brain showing the placement of the ECoG/EMG headmount and DBS electrodes. An image of the ECoG/EMG headmount and screws is shown in the bottom left. An image of the DBS electrodes is shown in the bottom right. **(b)** Video still from a sample sleep recording showing the experimental setup while a mouse is being recorded and stimulated. The ECoG/EMG preamplifier tether is shown in (1). The 360° commutator is shown in (2). The data acquisition system is shown in (3). All ECoG/EMG data is recorded with a PC computer and stored for analysis as shown in (4). The DBS stimulator box is shown in (5). The 360° commutator for the DBS tether is shown in (6). **(c)** A schematic illustration of the experimental timeline for recording sleep. **(d)** A schematic illustration of the DBS parameters. **(e)** A Nissl stain showing the bilateral placement of DBS electrodes between the interposed and fastigial cerebellar nuclei, and an illustration showing the relative anatomical locations of these structures in the brain. **(f)** Raw ECoG/EMG recordings from a control mouse, showing normal waveforms in the presence and absence of DBS during recording. **(g)** Quantification of the percentage of artifacts scored during sleep recordings. Points on **(g)** represent individual mice (n=9 per group). Source data and specific *p*-values for **g** are shown in Figure 1 – source data 1.

### Cerebellar nuclei DBS decreases total wake time and increases total REM and NREM time in *Ptf1a^Cre^;Vglut2^fx/fx^* and *Pdx1^Cre^;Vglut2^fx/fx^* mice

The relationship between sleep and motor function is crucial in dystonia. Observations have shown that after a good night’s sleep, motor symptoms become more manageable. This improvement is especially notable in the early morning after awakening^1,13^. However, existing therapies fail to improve sleep in patients^14^. Consequently, a primary objective of our study was to investigate how cerebellar nuclei DBS might improve overall sleep quality in *Ptf1a^Cre^;Vglut2^fx/fx^*and *Pdx1^Cre^;Vglut2^fx/fx^* mice. At baseline (Pre-DBS), representative hypnograms indicate that both *Ptf1a^Cre^;Vglut2^fx/fx^*and *Pdx1^Cre^;Vglut2^fx/fx^* mice display overall greater time awake and less time asleep relative to controls, similar to our previous findings^16^. Remarkably, during DBS, the proportions of wakefulness and sleep time began to converge across all mouse groups, demonstrating a rectification of sleep dysfunction. However, post-DBS ECoG recordings, when stimulation was withheld (Day-7), indicated that the sleep dysfunctions re-emerged in both *Ptf1a^Cre^;Vglut2^fx/fx^*and *Pdx1^Cre^;Vglut2^fx/fx^* mice (Figure 2A-C). We quantified the time spent in each state and found that indeed, DBS rectified the sleep deficits in the mutant mice. Both *Ptf1a^Cre^;Vglut2^fx/fx^*and *Pdx1^Cre^;Vglut2^fx/fx^* mice exhibited greater total sleep time, in response to cerebellar nuclei DBS. Notably, these changes demonstrated high temporal fidelity to DBS, as sleep-wake patterns reverted to baseline levels in the absence of stimulation (Figure 2D-F). We also found that DBS decreased the average length of wake bouts, increased the number of REM bouts, and increased the number of NREM bouts in *Ptf1a^Cre^;Vglut2^fx/fx^* and *Pdx1^Cre^;Vglut2^fx/fx^* mice (Figure 2G-I). Further, we calculated the latency to reach both REM and NREM sleep (Figure 2J). Our findings indicated that both groups of mutant mice, *Ptf1a^Cre^;Vglut2^fx/fx^* and *Pdx1^Cre^;Vglut2^fx/fx^*, experienced reduced latencies to REM and NREM sleep during DBS (Figure 2K, L). These findings indicate that cerebellar nuclei DBS effectively improves sleep timing and quality in both *Ptf1a^Cre^;Vglut2^fx/fx^* and *Pdx1^Cre^;Vglut2^fx/fx^* mice, with improvements closely tied to the DBS timing. Echoing our previous research^16^, these results point to cerebellar dysfunction, rather than motor impairment alone, as the likely origin of sleep deficits, given that both mutant mouse types, regardless of their motor condition, showed comparable sleep improvements during DBS.

**Figure 2:**
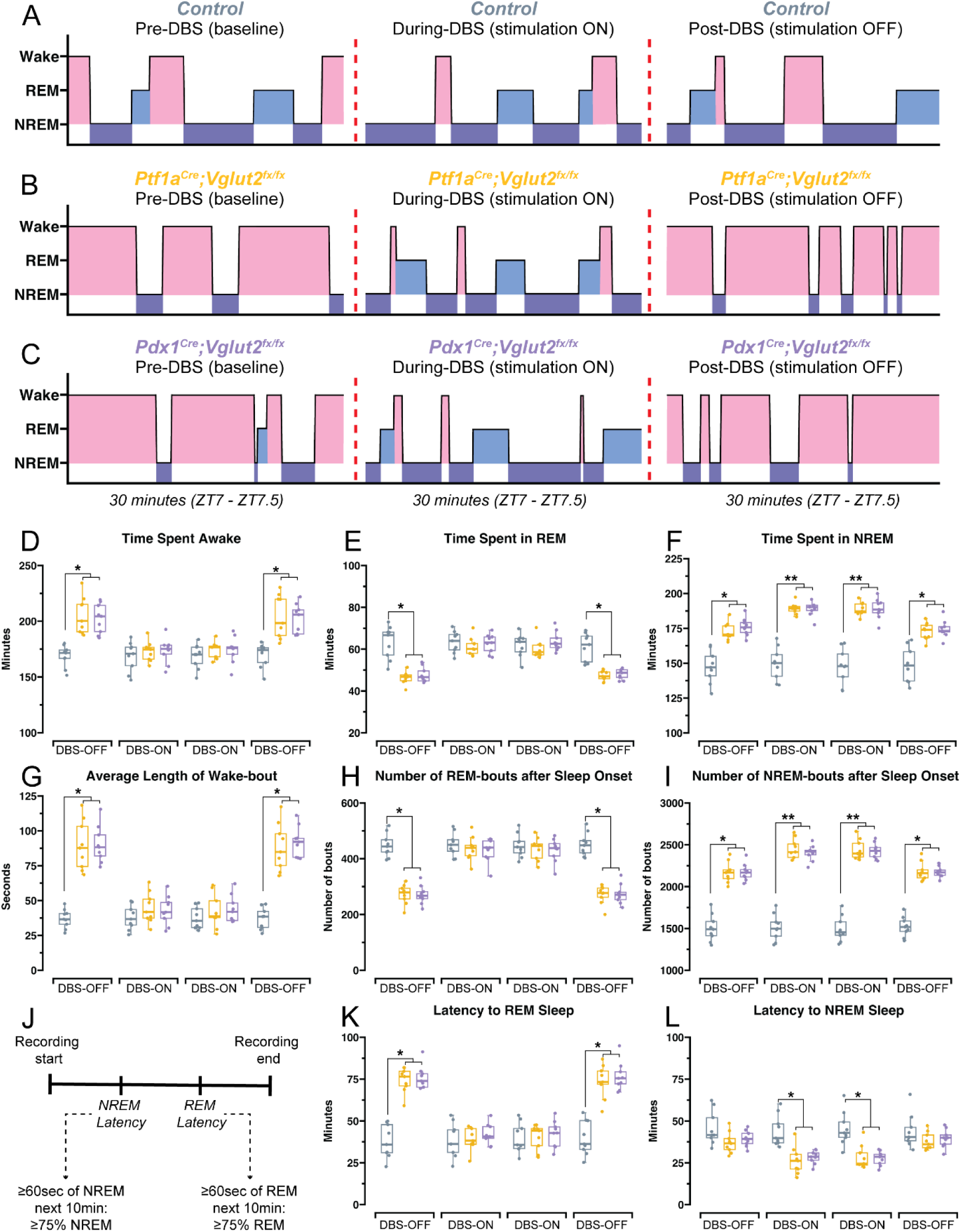
Cerebellar nuclei DBS impacts the same measures of sleep quality in both the *Ptf1a^Cre^;Vglut2^fx/fx^* and the *Pdx1^Cre^;Vglut2^fx/fx^* mutant mice. **(a)** Hypnograms taken from three different recording days, for a single representative control mouse for the same 30-minute period, ZT7-7.5, where ZT0 = lights ON, ZT1 = 1 hour after lights ON, etc. Periods of wake are highlighted in red, periods of REM are highlighted in light blue, periods of NREM are highlighted in dark purple. Dotted red lines separate different recording days, with different DBS status. **(b)** Same as **(a)** but for a *Ptf1a^Cre^;Vglut2^fx/fx^* mouse. **(c)** Same as **(a)** but for a *Pdx1^Cre^;Vglut2^fx/fx^* mouse. **(d)** Quantification of total time spent awake. Control mice in grey, *Ptf1a^Cre^;Vglut2^fx/fx^*mice in gold, *Pdx1^Cre^;Vglut2^fx/fx^* mice in purple. **(e)** Quantification of total time spent in REM. **(f)** Quantification of total time spent in NREM. **(g)** Quantification of average length of wake bouts. **(h)** Quantification of number of REM bouts after sleep onset. **(i)** Quantification of number of NREM bouts after sleep onset. **(j)** Schematic showing how NREM and REM latency were calculated, as in Hunsley & Palmiter, 2004^53^. **(k)** Quantification of latency to REM sleep. **(l)** Quantification of latency to NREM sleep. Points on **(d-i, k, l)** represent individual mice (n=9 per group). Source data and specific *p*-values for **d-i, k, l** is shown in Figure 2 – source data 1.

### Delta, beta, and gamma spectral frequency oscillations are modulated by cerebellar nuclei DBS

While cerebellar DBS has been established as an effective intervention for improving sleep in mouse models with dystonia, the underlying neurophysiological mechanisms remain an area of exploration. In particular, the modulation of spectral frequency oscillations during sleep provides a crucial window into understanding the intricacies of this therapeutic impact. Subsequently then, we sought to determine whether cerebellar nuclei DBS specifically influences these spectral frequency oscillations, which would shed light on its role in enhancing sleep quality. Arousal states are partially characterized by spectral frequency oscillations spanning a range from 0.5 to >100 Hz (Figure 3A). Thus, shifts between sleep stages can be marked by fluctuations in delta (0.5 Hz – 4 Hz in mice), theta (5 Hz – 8 Hz in mice), or alpha (8-13 Hz in mice) power, suggesting either an enhancement or deterioration in sleep quality^36,37^. Beta (13-30 Hz in mice) and gamma (35-44 Hz in mice) frequency bands can also act as indicators of sleep homeostasis, as they too tend to occur during specific arousal states (Figure 3B). We hypothesized that the existing differences in spectral power across the frequency bands of interest could be modulated alongside the patterns of sleep disruption during DBS. Upon analysis, we noted that indeed the baseline (pre-DBS) increase in frontal delta power in *Ptf1a^Cre^;Vglut2^fx/fx^*mice was significantly reduced during DBS (Figure 3C). Although the delta power in *Pdx1^Cre^;Vglut2^fx/fx^* mice was elevated during baseline, but not to the point of a significant difference, DBS did reduce delta power in these mice as well, matching the levels observed in *Ptf1a^Cre^;Vglut2^fx/fx^*mice and controls. Theta and alpha power remained unaltered for all mice across all conditions (Figure 3D, E). We also observed a reduction in pre-DBS beta and gamma power in the *Ptf1a^Cre^;Vglut2^fx/fx^*mice, which was normalized during DBS, as were the non-significant reductions in the *Pdx1^Cre^;Vglut2^fx/fx^*mice (Figure 3F, G). These data imply that the regulatory impacts of cerebellar nuclei DBS on neural circuitry extend beyond the cerebellum in this particular paradigm, evidenced by the modulation of the baseline changes in spectral frequency oscillations recorded in the *Ptf1a^Cre^;Vglut2^fx/fx^*and *Pdx1^Cre^;Vglut2^fx/fx^*mice.

**Figure 3:**
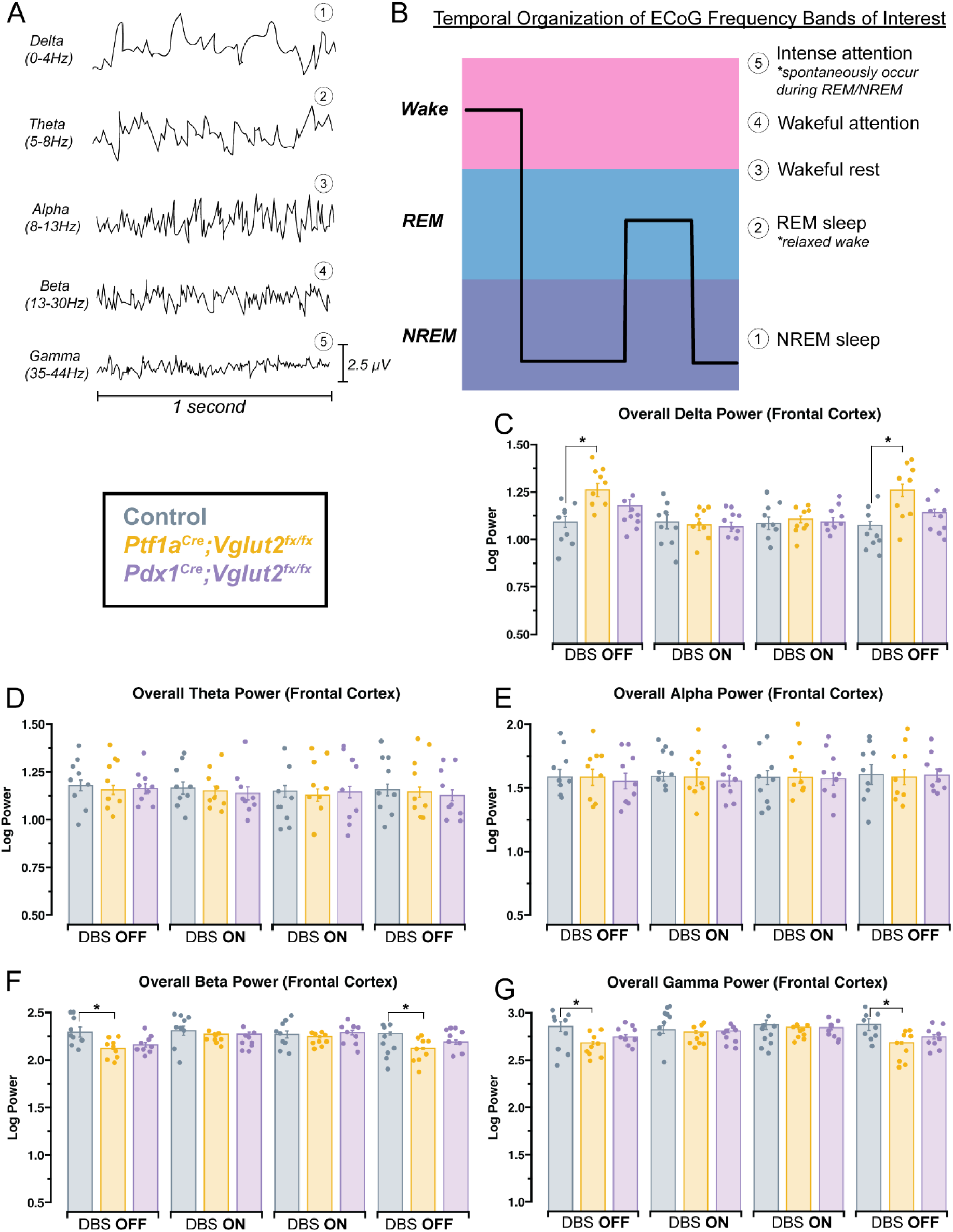
Cerebellar nuclei DBS normalizes differences in spectral frequency oscillations in the *Ptf1a^Cre^;Vglut2^fx/fx^* and the *Pdx1^Cre^;Vglut2^fx/fx^* mutant mice compared to control mice. **(a)** 1-second samples of raw ECoG waveforms for frequency bands of interest, from a control mouse. **(b)** Schematic illustration showing the relative arousal state in which different spectral frequency oscillations occur. **(c)** Quantification of delta power (0-4 Hz). Average power across the entire recording period. **(d)** Quantification of theta power (5-8 Hz). Average power across the entire recording period. **(e)** Quantification of alpha power (8-13 Hz). Average power across the entire recording period. **(f)** Quantification of beta power (13-30 Hz). Average power across the entire recording period. **(g)** Quantification of gamma power (35-44 Hz). Average power across the entire recording period. Points on **(c-g)** represent individual mice (n=9 per group). Source data and specific *p*-values for **c-g** are shown in Figure 3 – source data 1.

### The increased EMG power observed in *Ptf1a^Cre^;Vglut2^fx/fx^* mice is significantly reduced during cerebellar nuclei DBS, in all stages of sleep

The persistence of dystonia’s motor dysfunction during sleep is debated, with conflicting human studies suggesting both the resolution and possible continuation of these deficits impacting sleep quality^1,13,14^. Our previous study noted elevated trapezius EMG power in *Ptf1a^Cre^;Vglut2^fx/fx^*mice during both REM and NREM sleep, indicating ongoing motor symptoms^16^. We anticipated that cerebellar DBS, known for mitigating dystonic motor symptoms, would normalize these elevated EMG values. We recorded EMG signals using platinum-iridium wires that were surgically inserted into the trapezius muscles (Figure 4A), which we used to collect data continuously throughout the sleep recording period. By analyzing the power of the recorded raw trapezius muscle activity waveforms, specific to sleep stages before and during DBS, we showed that the stimulation normalized EMG activity. Specifically, the typically high-amplitude EMG waveforms in the *Ptf1a^Cre^;Vglut2^fx/fx^* mice during DBS shifted to more closely resemble those in control and *Pdx1^Cre^;Vglut2^fx/fx^*mice, both of which are devoid of motor dysfunction (Figure 4B). We calculated overall EMG power in the 0-30 Hz frequency band, previously used to diagnose dystonia in human patients^38^, and found that overall EMG activity was significantly reduced in the *Ptf1a^Cre^;Vglut2^fx/fx^*mice during DBS. Additionally, overall EMG power returned to baseline (elevated) levels in the absence of stimulation (Figure 4C). We further calculated EMG power only during periods of wake, REM, or NREM sleep and found that EMG power was similarly reduced in all states, in *Ptf1a^Cre^;Vglut2^fx/fx^*mice, with high temporal fidelity to DBS (Figure 4D-F). These findings confirm our previous work showing that, in the absence of DBS, dystonic muscle activity in *Ptf1a^Cre^;Vglut2^fx/fx^* mice persists in all sleep stages. Additionally, they suggest that our cerebellar nuclei DBS paradigm can move the elevated EMG power towards normalcy in *Ptf1a^Cre^;Vglut2^fx/fx^*mice, suggesting that correlates of the dystonic motor signatures are alleviated in all sleep states.

**Figure 4:**
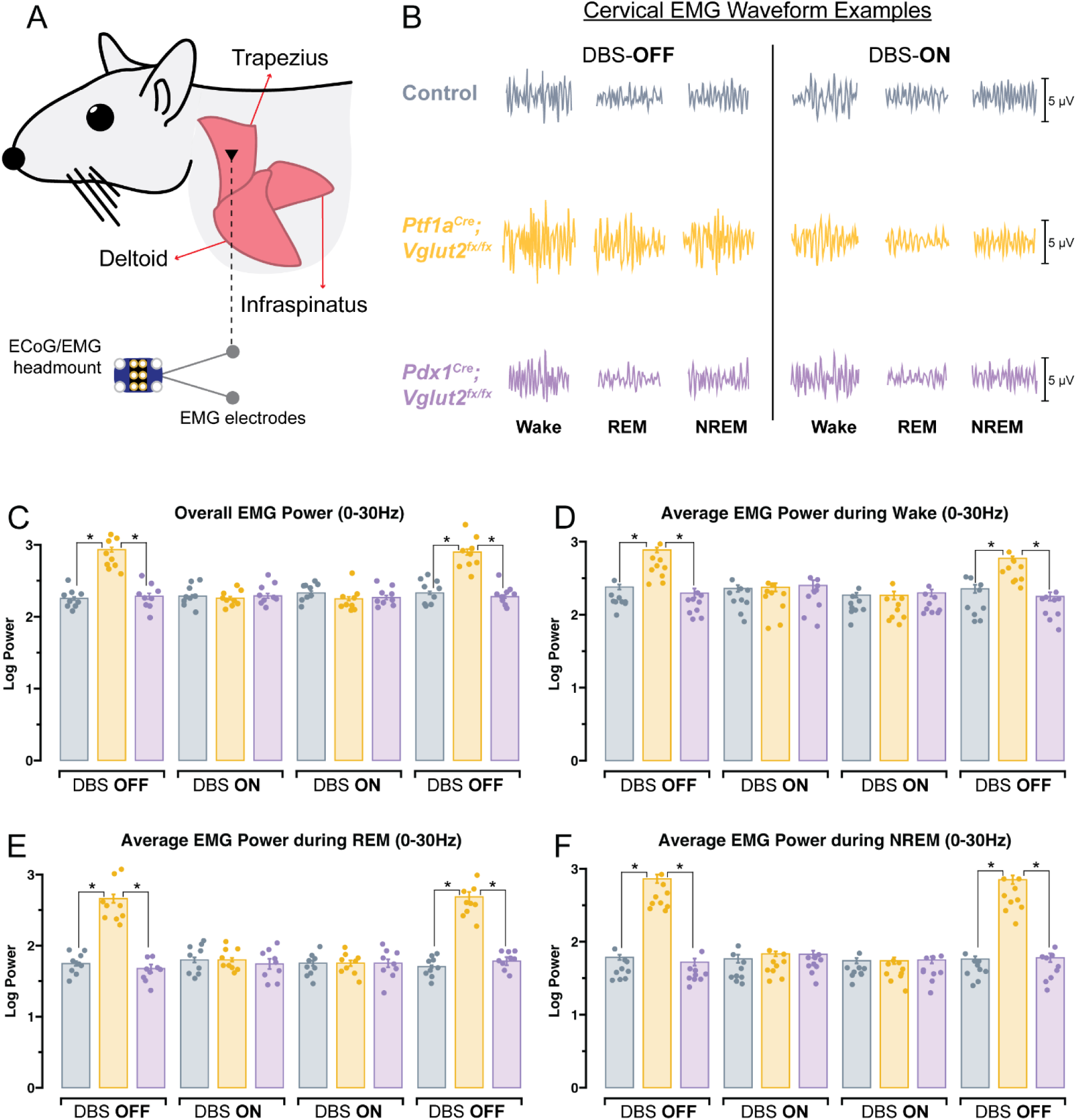
Cerebellar nuclei DBS reverses the abnormally increased EMG power of all sleep states observed in the *Ptf1a^Cre^;Vglut2^fx/fx^* mutant mice. **(a)** Schematic illustration of a mouse showing the musculature and relative placement of the EMG electrodes. **(b)** Raw EMG waveforms of trapezius activity for wake, REM and NREM sleep for mice of each group, across DBS-OFF and DBS-ON recording days. **(c)** Quantification of overall EMG activity (0-30 Hz). **(d)** Quantification of overall EMG activity only during periods of wake. **(e)** Quantification of overall EMG activity only during periods of REM sleep. **(f)** Quantification of overall EMG activity only during periods of NREM sleep. Points on **(c-f)** represent individual mice (n=9 per group). Source data and specific *p*-values for **c-f** are shown in Figure 4 – source data 1.

## Discussion

In this study, we examined both the motor and non-motor (sleep) symptoms of dystonia and the potential for therapeutic cerebellar DBS to improve sleep quality in mice. We dissected the role of the cerebellum, specifically the nuclei, as a target for DBS therapy in two mouse models of dystonia, *Ptf1a^Cre^;Vglut2^fx/fx^*and *Pdx1^Cre^;Vglut2^fx/fx^*, which have similar cerebellar deficits but differing motor phenotypes. Our results show that, while both mouse models demonstrate the same significant sleep deficits (particularly affecting REM sleep), the application of cerebellar nuclei DBS yielded significant improvements in both the quality and timing of sleep. During DBS, we observed a reduction in wakefulness and an increase in REM sleep in both mutant groups, indicating an overall reversal of these specific sleep deficits that are associated with dystonia. Moreover, we identified a notable decrease in the latency to REM and NREM sleep, addressing a prominent sleep dysfunction commonly reported in human dystonia patients^13,14^. Importantly, our findings suggest that the therapeutic potential of our cerebellar nuclei DBS paradigm extends beyond sleep regulation. An evaluation of motor symptoms in *Ptf1a^Cre^;Vglut2^fx/fx^* mice revealed an improvement across all sleep states, further demonstrating the potential efficacy of cerebellar nuclei DBS as a multi-dimensional intervention for dystonia. Beyond the modulation of key dystonia symptoms, this work also highlights DBS-induced modulation of the electrocorticogram (ECoG) patterns that define sleep states, providing additional insight into the neural underpinnings of sleep regulation, cerebellar function, and their intersection with motor dysfunctions.

While typical DBS therapies for motor disorders such as dystonia often target various subcortical structures including the basal ganglia and thalamus, recent evidence has suggested that cerebellar DBS shows promise for dystonia, which can be unresponsive to typical DBS targeting in a subpopulation of human patients^21,29^. In previous work, we demonstrated that the cerebellum, and in particular the interposed cerebellar nuclei, is an optimal target for DBS in mouse models for dystonia, tremor and ataxia^7,20,30,39^. As we sought to further develop cerebellar DBS as a multi-modal therapy to address sleep alongside motor deficits, we expanded our previous approach to include an additional mouse model (*Pdx1^Cre^;Vglut2^fx/fx^*) and also targeted the most medial of the cerebellar nuclei, the fastigial nucleus (Figure 1E). The fastigial nucleus was an attractive target for DBS given its role in non-motor functions, particularly sleep. The fastigial nucleus is known to have reciprocal projections with the hypothalamus, notably with the lateral hypothalamus, a region known to contribute extensively to regulating sleep-wake behavior^33^. Furthermore, of the cerebellar nuclei responses with sleep-dependent changes in activity, the trends for increased firing may be strongest in the fastigial^34,40^. While we have previously mentioned that the mechanisms through which DBS alleviates motor or non-motor symptoms remain unclear, in the case of the fastigial nucleus it may be that DBS improves sleep through the release of 5-hydroxyryptamine (serotonin). Indeed, it is known that electrical stimulation of the fastigial nucleus can modulate release of serotonin in the frontal lobe of rats^35^, and serotonin is known to play a complex role in sleep, affecting wakefulness, REM, and the ability to maintain sleep^41,42^. Additionally, it has been demonstrated that pharmacogenetic suppression of the 5HT-2A receptors in the fastigial nucleus is sufficient to mediate stress-induced dystonia in the *tottering* mouse^43^. Although an extensive analysis of the molecular mechanisms through which stimulation of the fastigial and interposed nuclei mediate improvements in sleep are outside the scope of this current study, the current data do provide a fruitful avenue for future research to improve or fine-tune therapeutic outcomes.

To stimulate the cerebellar region effectively, our targeting strategy was designed to encompass the space between the interposed and fastigial cerebellar nuclei. Notably, beyond the specific sleep-related connections of the fastigial nucleus, these circuits form connections with the vestibular nuclei, reticular formation, and thalamus, which collectively project many areas including the cerebral cortex. Such extensive connectivity underscores the cerebellum’s profound influence on both motor and non-motor functions. The combined evidence from our histological examination and the post-stimulation behavioral outcomes—namely, the reduction of motor dysfunction and sleep disturbances—indicates the successful targeting of both nuclei. Accordingly, the position of our stimulation electrodes was designed to span the distance between these nuclei. Given the conductive nature of white matter, we are confident that our electrical stimulation effectively reached both nuclei. However, we also recognize that by targeting the white matter situated between these nuclei, we are likely to also influence several cerebellar afferent and efferent pathways. This encompasses the mossy and climbing fiber inputs to the cerebellum, as well as internal axons passing over the cerebellar nuclei. Regardless, the potentially broad impact of DBS in this region could indeed be beneficial, aligning with our goal of modulating diverse behavioral modalities to address both sleep and motor deficits in our mouse models.

Given the roles of the interposed nucleus in regulating motor function, the fastigial nucleus in sleep-related processes, and the ability for DBS to affect global brain-states, we were particularly interested to know whether stimulation would negatively impact our ability to record sleep. Raw ECoG/EMG waveforms from a control mouse before and during cerebellar DBS suggested that typical waveform patterns were maintained in all cases, and that our ability to score sleep states was unaffected (Figure 1F). As non-biological electrical interference is known to act as a contributing factor to the occurrence of artifacts in ECoG recordings, we additionally quantified the percentage of scored artifacts in all mice, before, during, and after DBS. We found that DBS did not negatively affect the number of artifacts in the recorded data, as all mice consistently displayed <4% of scored artifacts across all recording days (Figure 1G).

Our initial objective was to determine if cerebellar DBS could enhance the overall sleep duration in *Ptf1a^Cre^;Vglut2^fx/fx^* and *Pdx1^Cre^;Vglut2^fx/fx^*mice. These mouse models, in line with our previous investigations, exhibit sleep irregularities akin to those seen in human patients, encompassing extended periods of wakefulness, diminished REM sleep, and prolonged latency to REM ^13,14,16^. Our analysis of total time spent in each state showed that during DBS, both groups demonstrated decreased total wake time and increased total REM time (Figure 2D, E). Interestingly, while wake and REM time was normalized by DBS (returned to control levels in both mutant groups), NREM time remained elevated in both *Ptf1a^Cre^;Vglut2^fx/fx^*and *Pdx1^Cre^;Vglut2^fx/fx^*mice, when DBS was delivered (Figure 2F). Previous work suggests that DBS delivered during sleep could extend the sleep stage in which it is administered^24^. Given that our DBS paradigm is continuous without specific temporal constraints, it raises the possibility that our stimulation indiscriminately increases all sleep states. However, in this case we would expect to see changes in sleep for control mice, which we did not. Another interpretation could be related to the “REM Rebound” phenomenon, where, following periods of sleep deprivation or significant stressors, both rodents and humans display extended and intensified REM sleep during the subsequent sleep period^44^. While it is uncertain whether a similar effect exists for NREM sleep, the observed increase in NREM time in our study could be attributed to a similar mechanism. Additionally, DBS may augment sleep pressure. Our findings do reveal a shortened latency to NREM sleep in both *Ptf1a^Cre^;Vglut2^fx/fx^*and *Pdx1^Cre^;Vglut2^fx/fx^*mice during DBS, suggesting that mice in both groups fall asleep more rapidly during stimulation (Figure 2L). However, if DBS universally increased sleep pressure, we would anticipate observing the same phenomenon in control mice, which is not the case. Alternatively, instead of enhancing sleep pressure, DBS may resolve an existing impediment to sleep pressure by correcting abnormal signaling in the cerebellar circuit of *Ptf1a^Cre^;Vglut2^fx/fx^* and *Pdx1^Cre^;Vglut2^fx/fx^* mice. In this case, we would anticipate DBS-induced changes in NREM latency to be specific to the mutant mice, while the controls would be unaffected. Such a mechanism would concur with the hypothesis that, at least in some conditions and in specific circuits, DBS could function by causing an “informational lesion” that impacts select synapses, ultimately correcting (or compensating for) convergent sites of dysfunction^28^.

Given that there are projections between the cerebellum and regions that not only regulate sleep, but specific stages of sleep (REM and NREM)^3,45,46^, we sought to determine if cerebellar DBS improved overall time spent sleeping via specific alterations in the length and frequency of individual wake, REM, or NREM bouts. Indeed, we previously found that the sleep deficits in the *Ptf1a^Cre^;Vglut2^fx/fx^*and *Pdx1^Cre^;Vglut2^fx/fx^* mice were driven by an increased wake bout length, a decreased number of REM bouts, and an increased number of NREM bouts. The same deficits were recapitulated in the current cohort of mice in the absence of DBS. During stimulation, we found that changes were induced in these same three measures of sleep architecture. For both *Ptf1a^Cre^;Vglut2^fx/fx^* and *Pdx1^Cre^;Vglut2^fx/fx^* mice during DBS, the average length of wake bouts was decreased, and the frequency of both REM and NREM bouts was increased (Figure 2G-I). The dynamics of these specific changes in sleep architecture do suggest, as we discussed previously, that cerebellar DBS in *Ptf1a^Cre^;Vglut2^fx/fx^*and *Pdx1^Cre^;Vglut2^fx/fx^*mice is either rectifying an impediment to natural sleep pressure or causing a dual REM-rebound and NREM-rebound effect, or both. Indeed, the combination of these two possible mechanisms of sleep rectification together would explain the increase of both REM and NREM frequency, even though only REM sleep is decreased in the mutant mice under baseline conditions. In final support of the dual sleep pressure and sleep rebound model, we also found that specific sleep latencies were normalized by cerebellar DBS in both *Ptf1a^Cre^;Vglut2^fx/fx^* and *Pdx1^Cre^;Vglut2^fx/fx^* mice. We have previously mentioned that latency to achieve NREM sleep was decreased by cerebellar DBS in both mutant groups. Notably however, we also found that REM latency was similarly decreased (Figure 2K, L). The rectification of REM latency is of particular interest, as achieving REM sleep is often a particular challenge faced by patients with dystonia^13,14^. Although the reduction in REM latency in the *Ptf1a^Cre^;Vglut2^fx/fx^* mouse model—characterized by overt dystonic motor dysfunction—could be ascribed to the alleviation of motor symptoms, the parallel reduction in latency observed in the *Pdx1^Cre^;Vglut2^fx/fx^* mice, which lack overt motor symptoms, implies a more fundamental role of cerebellar dysfunction, and cerebellar DBS. This points not only to the cerebellum’s integral involvement in these specific sleep impairments but also to the potential of cerebellum-focused therapeutics (invasive or non-invasive) for resolving different symptoms in neurological diseases.

Our ECoG spectral activity analysis presents noteworthy findings, which illuminate the fundamental contributors to the specific sleep deficits in *Ptf1a^Cre^;Vglut2^fx/fx^* and *Pdx1^Cre^;Vglut2^fx/fx^*mice. These findings demonstrate that the effects of our DBS are not limited to the cerebellum; they extend to and influence cortical activity. The detected decrease in delta power during DBS for *Ptf1a^Cre^;Vglut2^fx/fx^* mice (Figure 3C) concurs with independent studies that show a reduction in delta power across successive NREM sleep periods^47^. Moreover, the increase in the frequency of NREM bouts in the mutant mice during DBS (Figure 2I) parallels this observed decline in delta power. Sleep deprivation, known to trigger a surge in delta power^37^, is prominent when DBS is absent and the mutant mice exhibit severe sleep deficits and reduced sleep time. However, the implementation of DBS, by reducing wakefulness and increasing sleep duration (Figure 2D-F), could mitigate the state of sleep deprivation, leading to the anticipated decrease in delta power during stimulation. Our observations of enhanced beta and gamma power during DBS add further depth to our understanding of how cerebellar DBS influences sleep in the *Ptf1a^Cre^;Vglut2^fx/fx^* and *Pdx1^Cre^;Vglut2^fx/fx^* mice. It is known that diminished beta power is linked with suboptimal sleep quality and could potentially suggest obstructive sleep apnea^48^. Even though the presence of obstructive sleep apnea in the *Ptf1a^Cre^;Vglut2^fx/fx^* mice has not been determined, we interpret the increase in beta power during DBS (Figure 3F) as possibly indicative of a restoration of motor function, a change we know our stimulation paradigm can induce. Meanwhile, the rise in gamma power during DBS (Figure 3G) could stem from an overall increase in REM and NREM sleep durations. Given the known spontaneous occurrence of gamma oscillations during both REM and NREM sleep ^49^, the overall increase in gamma power could correspond to the lengthened REM and NREM periods during DBS for the mutant mice. Lastly, we observed that *Pdx1^Cre^;Vglut2^fx/fx^* mice did not exhibit significant changes in delta, beta, or gamma power in the absence of DBS and that power in these bands does not differ significantly from that of *Ptf1a^Cre^;Vglut2^fx/fx^*mice or controls (Figure 3C, F, G). We have previously speculated that this may be due to their ‘intermediate’ cerebellar and motor phenotype, arising from the different pattern of olivocerebellar VGLUT2 deletion relative to the *Ptf1a^Cre^;Vglut2^fx/fx^*mice^16^. In this case, the observed spectral differences between *Ptf1a^Cre^;Vglut2^fx/fx^* and *Pdx1^Cre^;Vglut2^fx/fx^* mice may serve as biomarkers of disease severity, as shifts in spectral power can vary in direction and magnitude across different diseases, even when the affected individuals all exhibit similarly disrupted sleep^37,50^. We do also recognize that spectral differences in sleep between *Ptf1a^Cre^;Vglut2^fx/fx^* and *Pdx1^Cre^;Vglut2^fx/fx^* mice may intertwine with alterations in the motor program, considering the known influence of movement patterns on ECoG spectral activity^51^. Nevertheless, despite their “intermediate” spectral power differences, during DBS, the *Pdx1^Cre^;Vglut2^fx/fx^* mice demonstrate changes in spectral power that align directionally with those observed in the *Ptf1a^Cre^;Vglut2^fx/fx^*mice, reinforcing the robustness of the impact of cerebellar DBS in disease circuits across genetic models of dystonia.

The mechanistic implications of our EMG results bear special consideration from a therapeutic perspective. Traditional approaches, such as botulinum toxin injections, and even traditional DBS have demonstrated effectiveness in addressing the motor symptoms of dystonia. However, these methods have typically fallen short in simultaneously addressing the non-motor (sleep) symptoms of the disorder^14^. Our study reveals that our cerebellar DBS paradigm can potentially address both facets. Raw EMG recordings from the trapezius muscle of the *Ptf1a^Cre^;Vglut2^fx/fx^* mice corroborate the pervasive motor dysfunction across all arousal states, as indicated by the high-amplitude waveform patterns characteristic of dystonic motor contractions (Figure 4A, B). Furthermore, our analysis within the 0-30 Hz frequency band of EMG power showed a marked reduction not only overall during DBS (Figure 4C), but also within specific arousal states of wake, REM, or NREM sleep (Figure 4D-F). The result of DBS decreasing EMG power during REM sleep is particularly striking, given that muscle atonia, typically a defining characteristic of this sleep state, was not visible in the *Ptf1a^Cre^;Vglut2^fx/fx^*mice without the aid of DBS. This finding indicates that our DBS paradigm, targeting the fastigial and interposed cerebellar nuclei, can address motor dysfunction across arousal states, including sleep - an improvement on our previously tested paradigms targeting only the interposed nuclei^7,30^. When considered in tandem with our earlier findings showing the beneficial impact of cerebellar DBS on sleep dysfunction in both the *Ptf1a^Cre^;Vglut2^fx/fx^*and the *Pdx1^Cre^;Vglut2^fx/fx^*mutant mice, this data underscores the potential of our DBS paradigm as a multi-functional therapeutic strategy.

Collectively, our findings highlight the promising therapeutic potential of cerebellar DBS in dystonia, demonstrating its capacity to simultaneously reduce both the motor and non-motor symptoms that can exacerbate one another. This is particularly noteworthy as the importance of addressing sleep dysfunction is increasingly recognized, not only in dystonia but in diverse societal contexts, given its substantial impact on quality of life, propensity to lead to various comorbidities, and association with impaired motor function and learning. These factors inspire a compelling model where sleep and motor dysfunctions in dystonia form a self-perpetuating cycle, in which motor dysfunction contributes to sleep impairments, and sleep impairments in turn result in worsened patterns of synaptic renormalization, motor learning, and motor dysfunction. Therefore, the dual-effect DBS strategy we have tested signifies a considerable advancement in the therapeutic approach towards dystonia, offering a promising avenue for future research and clinical application. Our results further reinforce the central role of the cerebellum within broader neural circuits, and consequently, the profound potential of cerebellar-targeted interventions. The broad-reaching effects of cerebellar dysfunction, and the correspondingly widespread implications of its modulation through DBS, underscore the necessity and promise of cerebellar interventions in the context of motor disease and beyond. Therefore, our cerebellar DBS paradigm stands to serve as a comprehensive potential platform for addressing the spectrum of dystonia symptoms.

## Ethics

Animal experimentation: All animals (mice) were housed in an AALAS-certified facility that operates on a 14-hour light cycle. Husbandry, housing, euthanasia, and experimental guidelines were reviewed and approved by the Institutional Animal Care and Use Committee (IACUC) of Baylor College of Medicine (protocol number: AN-5996).

## Methods

### Animals

All mice used in this study were housed in a Level 3, AALAS-certified facility. All experiments and studies that involved mice were reviewed and approved by the Institutional Animal Care and Use Committee of Baylor College of Medicine (BCM AN-5996). Dr. Chris Wright (Vanderbilt University School of Medicine) kindly provided the *Ptf1a^Cre^* mice. We purchased the *Pdx1^Cre^* (*Pdx-Cre*, #014647) and *Vglut2^floxed^* (*Vglut2^fx^*, #012898) mice from The Jackson Laboratory (Bar Harbor, ME, USA) and then maintained them in our colony using a standard breeding scheme. The conditional knock-out mice that resulted in dystonia were generated by crossing *Ptf1a^Cre^;Vglut2^fx/+^* heterozygote mice or *Pdx1^Cre^;Vglut2^fx/+^* heterozygote mice with homozygote *Vglut2^fx/fx^* mice. *Pdx1^Cre^;Vglut2^fx/fx^* and *Ptf1a^Cre^;Vglut2^fx/fx^* mice were considered experimental animals. A full description of the genotyping details (e.g., primer sequences and the use of a standard polymerase chain reaction) and phenotype for the *Ptf1a^Cre^;Vglut2^fx/fx^* mouse was provided in White and Sillitoe, 2017^7^. A full description of the genotype and the initial observations of the phenotype of the *Pdx1^Cre^;Vglut2^fx/fx^* mouse was provided in Lackey, 2022^8^. All littermates lacking Cre upon genotyping were considered control mice. Ear punches were collected before weaning and used for genotyping and identification of the different alleles. For all experiments, we bred mice using standard timed pregnancies, noon on the day a vaginal plug was detected was considered embryonic day (E)0.5 and postnatal day (P)0 was defined as the day of birth. Mice of both sexes were used, and the data combined in all experiments.

### Tissue preparation and Histology

For perfusion fixation, animals were deeply anesthetized with 2, 2, 2-tribromoethanol (Avertin), and then perfused through the heart with 0.1 m PBS (pH 7.2), followed by 4% paraformaldehyde (PFA) diluted in PBS. The brains from the perfused mice were postfixed for 24–48 h in 4% PFA and then cryoprotected stepwise in PBS-buffered sucrose solutions (15 and 30% each time until the brain sunk). Serial 40-μm-thick coronal tissue sections were cut on a cryostat, and then each tissue section collected in PBS and processed free floating.

Nissl staining was performed by first mounting the tissue sections on slides and allowing them to dry on the slides overnight. Mounted sections were immersed in 100% xylene two times for 5 min and then put through a rehydration series which consisted of 3 immersions in 100% ethanol followed by 95% ethanol, 70% ethanol, 50% ethanol and then tap water, for two minutes per step. The tissue sections were then immersed in cresyl violet solution for ∼30 seconds or until the stain was sufficiently dark. The tissue sections were then dehydrated using the reversed order of the rehydration series (described above) followed by xylene, with 30s per step. Cytoseal XYL mounting media (Thermo Scientific, Waltham, MA, USA, #22-050-262) and a coverslip was then immediately placed on the slides.

### Surgical procedures

Prior to surgery, mice were given preemptive analgesics (buprenorphine, 1 mg/kg subcutaneous (SC), and meloxicam, 5 mg/kg SC) with continued application as part of the standard 3 day post-operative procedure. Mice were anesthetized with isoflurane and placed into a stereotaxic device, which continued to deliver isoflurane throughout the surgery. Each mouse was implanted with a prefabricated ECoG/EMG headmount (Pinnacle Technology, Lawrence KS, #8201) with 0.10” EEG screws to secure headmounts to the skull (Pinnacle Technology, Lawrence KS, #8209). A midline incision was made, and the skull was exposed. The headmount was affixed to the skull using cyanoacrylate glue to hold in place while pilot holes for screws were made and screws were inserted. Screws were placed bilaterally over the parietal cortex and frontal cortex. A small amount of silver epoxy (Pinnacle Technology, Lawrence KS, #8226) was applied to the screw-headmount connection. Platinum-iridium EMG wires on the prefabricated headmount were placed under the skin of the neck, resting directly on the trapezius muscles.

During the same surgery to implant ECoG/EMG headmounts, two small craniotomies were performed bilaterally and dorsal to the region between the fastigial and interposed cerebellar nuclei, through which two custom 50 mm twisted bipolar Tungsten DBS electrodes were lowered just into the region of interest (PlasticsOne, Roanoke, VA, USA; #8IMS303T3B01).

The headmount and DBS electrodes were permanently affixed to the skull using ‘Cold-Cure’ dental cement (A-M systems, #525000 and #526000). Mice were allowed to recover for 3-4 days before being fitted with a preamplifier (Pinnacle Technology, Lawrence KS, #8202) and tethered to the recording device (Pinnacle Technology, Lawrence KS, #8204 and #8206-HR) and the deep brain stimulation device (Multi Channel Systems, #STG4002).

### ECoG/EMG sleep recording

Mice were recorded in light and temperature-controlled rooms, for 8-hours, at the same time of day for every mouse and for each day. The first hour of recording was considered the acclimation period and was therefore excluded from final analysis. Food and water were available *ad libitum* throughout the recording days. Mice were tethered to both the sleep recording hardware and the DBS stimulator for each day of recording. The DBS stimulator was turned on for every day of recording, but stimulation was delivered only on days labeled “DBS ON”.

### Sleep scoring and analysis of sleep data

Sleep stages and artifacts were automatically scored offline with SPINDLE ^52^, as previously described ^16^. For spectral frequency analysis of ECoG and EMG activity, raw files were also pre-processed in MATLAB (MathWorks) using the free toolkit EEGLAB (UC San Diego). Scored files were downloaded from SPINDLE as a .csv and statistical analysis was performed in R v4.1.2.

### Data analysis and statistics

Data are presented as mean ± SEM and analyzed as a mixed ANOVA followed by post-hoc pairwise comparisons with Tukey’s method for multiple comparisons correction. *p* < 0.05 was considered as statistically significant. All statistical analyses were performed using R v4.1.2.

## Data availability

All data generated or analyzed in this study are included in the manuscript and supporting files.

## Acknowledgements and Funding Sources

This work was supported by Baylor College of Medicine (BCM), Texas Children’s Hospital, The Hamill Foundation, and the National Institutes of Neurological Disorders and Stroke (NINDS) R01NS100874, R01NS119301, and R01NS127435 to RVS. Research reported in this publication was supported by the Eunice Kennedy Shriver National Institute of Child Health & Human Development of the National Institutes of Health under Award Number P50HD103555 for use of the Animal Behavior Core and the Cell and Tissue Pathogenesis Core (BCM IDDRC). The content is solely the responsibility of the authors and does not necessarily represent the official views of the National Institutes of Health. Support was also provided by a Dystonia Medical Research Foundation (DMRF) grant to RVS and a DMRF fellowship to LHK.

## Author contributions

Technical and conceptual ideas in this work were conceived by LESL and RVS. LESL performed experiments. LESL, LHK, and RVS performed data analysis and data interpretation. LESL, LHK and RVS wrote the manuscript. LESL, LHK, and RVS edited the manuscript.

## References

1. Smith, B. Lived Experience with Dystonia. 633 https://dystoniasurveys.files.wordpress.com/2021/08/lived-experience-dystonia-patient-surveys-2020-2021.pdf (2021).

2. Balint, B. et al. Dystonia. Nat. Rev. Dis. Primer 4, 1–23 (2018).

3. Salazar Leon, L. E. & Sillitoe, R. V. Potential interactions between cerebellar dysfunction and sleep disturbances in dystonia. Dystonia 1, 10691 (2022).

4. Corp, D. T. et al. Network localization of cervical dystonia based on causal brain lesions. Brain 142, 1660–1674 (2019).

5. Hendrix, C. M. & Vitek, J. L. Toward a network model of dystonia. Ann. N. Y. Acad. Sci. 1265, 46–55 (2012).

6. Fremont, R., Tewari, A., Angueyra, C. & Khodakhah, K. A role for cerebellum in the hereditary dystonia DYT1. eLife 6, e22775 (2017).

7. White, J. J. & Sillitoe, R. V. Genetic silencing of olivocerebellar synapses causes dystonia-like behaviour in mice. Nat. Commun. 8, 14912 (2017).

8. Elizabeth Lackey. Neural function of Cerebellar inputs in Dystonia-like behavior. (Baylor College of Medicine, 2022).

9. Sadnicka, A., Hoffland, B. S., Bhatia, K. P., van de Warrenburg, B. P. & Edwards, M. J. The cerebellum in dystonia - help or hindrance? Clin. Neurophysiol. Off. J. Int. Fed. Clin. Neurophysiol. 123, 65–70 (2012).

10. Aïssa, H. B. et al. Functional abnormalities in the cerebello-thalamic pathways in a mouse model of DYT25 dystonia. eLife 11, e79135 (2022).

11. Canto, C. B., Onuki, Y., Bruinsma, B., Werf, Y. D. van der & Zeeuw, C. I. D. The Sleeping Cerebellum. Trends Neurosci. 40, 309–323 (2017).

12. Cunchillos, J. D. & De Andrés, I. Participation of the cerebellum in the regulation of the sleep-wakefulness cycle. Results in cerebellectomized cats. Electroencephalogr. Clin. Neurophysiol. 53, 549–558 (1982).

13. Antelmi, E. et al. Modulation of the Muscle Activity During Sleep in Cervical Dystonia. Sleep 40, zsx088 (2017).

14. Eichenseer, S. R., Stebbins, G. T. & Comella, C. L. Beyond a motor disorder: A prospective evaluation of sleep quality in cervical dystonia. Parkinsonism Relat. Disord. 20, 405–408 (2014).

15. Hertenstein, E. et al. Sleep in patients with primary dystonia: A systematic review on the state of research and perspectives. Sleep Med. Rev. 26, 95–107 (2016).

16. Leon, L. E. S. & Sillitoe, R. V. Disrupted sleep in dystonia depends on cerebellar function but not motor symptoms in mice. 2023.02.09.527916 Preprint at 10.1101/2023.02.09.527916 (2023).

17. Boudreau, P., Dumont, G. A. & Boivin, D. B. Circadian adaptation to night shift work influences sleep, performance, mood and the autonomic modulation of the heart. PloS One 8, e70813 (2013).

18. James, S. M., Honn, K. A., Gaddameedhi, S. & Van Dongen, H. P. A. Shift Work: Disrupted Circadian Rhythms and Sleep—Implications for Health and Well-Being. Curr. Sleep Med. Rep. 3, 104–112 (2017).

19. Smit, M. et al. Fatigue, Sleep Disturbances, and Their Influence on Quality of Life in Cervical Dystonia Patients. Mov. Disord. Clin. Pract. 4, 517–523 (2017).

20. Brown, E. G. et al. Cerebellar Deep Brain Stimulation for Acquired Hemidystonia. Mov. Disord. Clin. Pract. 7, 188–193 (2020).

21. Krauss, J. K., Yianni, J., Loher, T. J. & Aziz, T. Z. Deep brain stimulation for dystonia. J. Clin. Neurophysiol. Off. Publ. Am. Electroencephalogr. Soc. 21, 18–30 (2004).

22. Wichmann, T. & DeLong, M. R. Deep-Brain Stimulation for Basal Ganglia Disorders. Basal Ganglia 1, 65–77 (2011).

23. Horisawa, S. et al. Intermittent Ultralow-Frequency Low-Amplitude Deep Cerebellar Stimulation for Movement Disorders. Mov. Disord. Clin. Pract. **n/a**,.

24. Amara, A. W., Watts, R. L. & Walker, H. C. The effects of deep brain stimulation on sleep in Parkinson’s disease. Ther. Adv. Neurol. Disord. 4, 15–24 (2011).

25. Bjerknes, S., Skogseid, I. M., Hauge, T. J., Dietrichs, E. & Toft, M. Subthalamic deep brain stimulation improves sleep and excessive sweating in Parkinson’s disease. Npj Park. Dis. 6, 1–7 (2020).

26. Zuzuárregui, J. R. P. & Ostrem, J. L. The Impact of Deep Brain Stimulation on Sleep in Parkinson’s Disease: An update. J. Park. Dis. 10, 393–404 (2020).

27. Gilron, R. et al. Sleep-Aware Adaptive Deep Brain Stimulation Control: Chronic Use at Home With Dual Independent Linear Discriminate Detectors. Front. Neurosci. 15, (2021).

28. Lozano, A. M. & Eltahawy, H. How does DBS work? Suppl. Clin. Neurophysiol. 57, 733–736 (2004).

29. Tai, C.-H. & Tseng, S.-H. Cerebellar deep brain stimulation for movement disorders. Neurobiol. Dis. 175, 105899 (2022).

30. Miterko, L. N. et al. Neuromodulation of the cerebellum rescues movement in a mouse model of ataxia. Nat. Commun. 12, 1295 (2021).

31. Mencacci, N. E. et al. Biallelic variants in *TSPOAP1*, encoding the active-zone protein RIMBP1, cause autosomal recessive dystonia. J. Clin. Invest. 131, (2021).

32. Tarnecki, R. Functional connections between neurons of interpositus nucleus of the cerebellum and the red nucleus. Behav. Brain Res. 28, 117–125 (1988).

33. Zhang, X.-Y., Wang, J.-J. & Zhu, J.-N. Cerebellar fastigial nucleus: from anatomic construction to physiological functions. Cerebellum Ataxias 3, 9 (2016).

34. Palmer, C. Interpositus and fastigial unit activity during sleep and waking in the cat. Electroencephalogr. Clin. Neurophysiol. 46, 357–370 (1979).

35. Wang, J., Dong, W.-W., Zhang, W.-H., Zheng, J. & Wang, X. Electrical stimulation of cerebellar fastigial nucleus: mechanism of neuroprotection and prospects for clinical application against cerebral ischemia. CNS Neurosci. Ther. 20, 710–716 (2014).

36. Bjorness, T. E., Booth, V. & Poe, G. R. Hippocampal theta power pressure builds over non-REM sleep and dissipates within REM sleep episodes. Arch. Ital. Biol. 156, 112–126 (2018).

37. Long, S. et al. Sleep Quality and Electroencephalogram Delta Power. Front. Neurosci. 15, (2021).

38. Go, S. A., Coleman-Wood, K. & Kaufman, K. R. Frequency analysis of lower extremity electromyography signals for the quantitative diagnosis of dystonia. J. Electromyogr. Kinesiol. Off. J. Int. Soc. Electrophysiol. Kinesiol. 24, 31–36 (2014).

39. Brown, A. M. et al. Purkinje cell misfiring generates high-amplitude action tremors that are corrected by cerebellar deep brain stimulation. eLife 9, e51928 (2020).

40. Hobson, J. A. & McCarley, R. W. Spontaneous discharge rates of cat cerebellar Purkinje cells in sleep and waking. Electroencephalogr. Clin. Neurophysiol. 33, 457–469 (1972).

41. Portas, C. M., Bjorvatn, B. & Ursin, R. Serotonin and the sleep/wake cycle: special emphasis on microdialysis studies. Prog. Neurobiol. 60, 13–35 (2000).

42. Whitney, M. S. et al. Adult Brain Serotonin Deficiency Causes Hyperactivity, Circadian Disruption, and Elimination of Siestas. J. Neurosci. Off. J. Soc. Neurosci. 36, 9828–9842 (2016).

43. Kim, J. E. et al. Cerebellar 5HT-2A receptor mediates stress-induced onset of dystonia. Sci. Adv. (2021) doi:10.1126/sciadv.abb5735.

44. Feriante, J. & Singh, S. REM Rebound Effect. in StatPearls (StatPearls Publishing, 2023).

45. Van Dort, C. J. et al. Optogenetic activation of cholinergic neurons in the PPT or LDT induces REM sleep. Proc. Natl. Acad. Sci. U. S. A. 112, 584–589 (2015).

46. Eban-Rothschild, A., Rothschild, G., Giardino, W. J., Jones, J. R. & de Lecea, L. VTA dopaminergic neurons regulate ethologically relevant sleep–wake behaviors. Nat. Neurosci. 19, 1356–1366 (2016).

47. Darchia, N., Campbell, I. G., Tan, X. & Feinberg, I. Kinetics of NREM Delta EEG Power Density Across NREM Periods Depend on Age and on Delta-Band Designation. Sleep 30, 71–79 (2007).

48. Liu, S. et al. EEG Power Spectral Analysis of Abnormal Cortical Activations During REM/NREM Sleep in Obstructive Sleep Apnea. Front. Neurol. 12, (2021).

49. Valderrama, M. et al. Human Gamma Oscillations during Slow Wave Sleep. PLoS ONE 7, e33477 (2012).

50. Davis, C. J., Clinton, J. M., Jewett, K. A., Zielinski, M. R. & Krueger, J. M. Delta Wave Power: An Independent Sleep Phenotype or Epiphenomenon? J. Clin. Sleep Med. JCSM Off. Publ. Am. Acad. Sleep Med. 7, S16–S18 (2011).

51. Mihajlovic, V., Patki, S. & Grundlehner, B. The impact of head movements on EEG and contact impedance: an adaptive filtering solution for motion artifact reduction. Annu. Int. Conf. IEEE Eng. Med. Biol. Soc. IEEE Eng. Med. Biol. Soc. Annu. Int. Conf. 2014, 5064–5067 (2014).

52. Miladinović, Đ., et al. SPINDLE: End-to-end learning from EEG/EMG to extrapolate animal sleep scoring across experimental settings, labs and species. PLOS Comput. Biol. 15, e1006968 (2019).

53. Hunsley, M. S. & Palmiter, R. D. Altered sleep latency and arousal regulation in mice lacking norepinephrine. Pharmacol. Biochem. Behav. 78, 765–773 (2004).

